# A putative human infertility allele of the meiotic recombinase DMC1 does not affect fertility in mice

**DOI:** 10.1101/313890

**Authors:** Tina N. Tran, John C. Schimenti

## Abstract

Whole exome or genome sequencing is becoming routine in clinical situations for identifying mutations underlying presumed genetic causes of disease, including infertility. While this is a powerful approach for implicating polymorphisms or de novo mutations in genes plausibly related to the phenotype, a greater challenge is to definitively prove causality. This is a crucial requisite for treatment, especially for infertility, in which validation options are limited. In this study, we created a mouse model of a putative infertility allele, *DMC1^M200V^*. *DMC1* encodes a RecA homolog essential for meiotic recombination and fertility in mice. This allele was originally implicated as being responsible for sterility of a homozygous African woman, a conclusion supported by subsequent biochemical analyses of the mutant protein and by studies of yeast with the orthologous amino acid change. Here, we found that *Dmc1^M200V/M200V^* male and female mice are fully fertile and do not exhibit any gonadal abnormalities. Detailed immunocytological analysis of meiosis revealed no defects suggestive of compromised fertility. This study serves as a cautionary tale for making conclusions about consequences of genetic variants, especially with respect to infertility, and emphasizes the importance of conducting relevant biological assays for making accurate diagnoses in the era of genomic medicine.

## INTRODUCTION

Surveys estimate that 10-15% of the United States population experiences some form of infertility (Chandra, 2013), and it is believed that as many as half of infertility cases have an underlying genetic basis. Over the last decade, candidate gene sequencing and whole exome/genome re-sequencing have become increasingly used clinically for identifying potential causes of presumed genetic disorders, including infertility. Several studies have employed the general approach of identifying a candidate variant/mutation in a sterile patient(s), sequencing the suspect gene in a number of normal patients, and then suggesting causation of the candidate mutation if it is unique (or homozygous in) only the infertile people (Sato et al. 2006; Miyamoto et al. 2008; Miyamoto et al. 2009; França et al. 2017) However, such correlation is not proof, as illustrated in the case of a purported globozoospermia allele of *Spata16* (Dam et al. 2007) that was shown by mouse modeling to not impact fertility (Fujihara et al. 2017).

A major strategy for identifying and implicating putative disease-mutations is the use of algorithms designed to predict the impact of mutations on protein function. Popular algorithms such as Polyphen-2, SIFT, and FATHMM utilize direct or machine-learning strategies to assess protein sequence, structure, and amino acid properties to generate predictions (Kumar et al. 2009; Adzhubei et al. 2010; Shihab et al. 2013). Nonetheless, recent work has found that the predictions of damaging alleles are frequently inaccurate, and many predicted null mutations cause subtle deficiencies at best (Wang et al. 2018; Miosge et al. 2015). In addition, studies specifically aimed at fertility effects *in vivo* found that as low as 25% of predicted high-damaging variants cause any detectable phenotype (Singh and Schimenti 2015). Although computational approaches can be useful, these studies emphasize that current algorithms may not be sufficiently reliable for high-confidence assessment of the physiological impact of variants, especially in a clinical context.

In this present study, we took a closer look at a variant of human *DMC1*, a methionine-to-valine change at amino acid position 200 (*Dmc1^M200V^*). *Dmc1* is a meiosis-specific homolog of *E. coli RecA* and is essential for the repair of SPO11/TOP6BL-induced double strand breaks (DSBs) by homologous recombination (Yoshida et al. 1998; Bishop et al. 1992; Robert et al. 2016; Keeney et al. 1997), a process that is essential for pairing of chromosome homologs. *Dmc1* knockout mice are sterile due to arrest of meiocytes stemming from failed homologous chromosome synapsis (Yoshida et al. 1998; D. Pittman et al. 1998) and activation of the meiotic DNA damage checkpoint (Bolcun-Filas et al. 2014). *DMC1^M200V^* was implicated as being responsible for premature ovarian failure in an African woman who was homozygous for this allele (Mandon-Pépin et al. 2008; Hikiba et al. 2008). Follow-up studies of this allele including crystallographic characterization, enzymatic assays, and phenotypic analysis of yeast bearing the orthologous amino acid change, demonstrated impaired function of the protein (Hikiba et al. 2008). We sought to expand upon these findings by introducing the M200V allele in the mouse ortholog of *Dmc1.* Here, we report that *Dmc1^M200V/M200V^* male and female mice are fertile, have normal fecundity and gamete numbers, and have no meiotic defects. Our findings underscore the potential difficulties in reliably assessing allele pathogenicity and argue for the use of relevant *in vivo* functional assays to guide therapeutic actions.

## RESULTS

### *Dmc1^M200V/M200V^* mice are fertile and phenotypically similar to wild type

To generate mice modeling the human *DMC1^M200V^* allele, CRISPR/Cas9-mediated genome editing in single-celled zygotes was performed. As diagrammed in Figure 1A and B, a single-stranded oligodeoxynucleotide (ssODN) was used as a template to introduce two nucleotide changes into codon 200 via homologous recombination, causing this codon to encode valine instead of methionine. Founder mice with the desired mutation were identified (Fig. 1C), backcrossed into FVB/NJ for at least two generations, then intercrossed to generate heterozygous and homozygous mice for phenotypic analysis.

**Figure 1.**
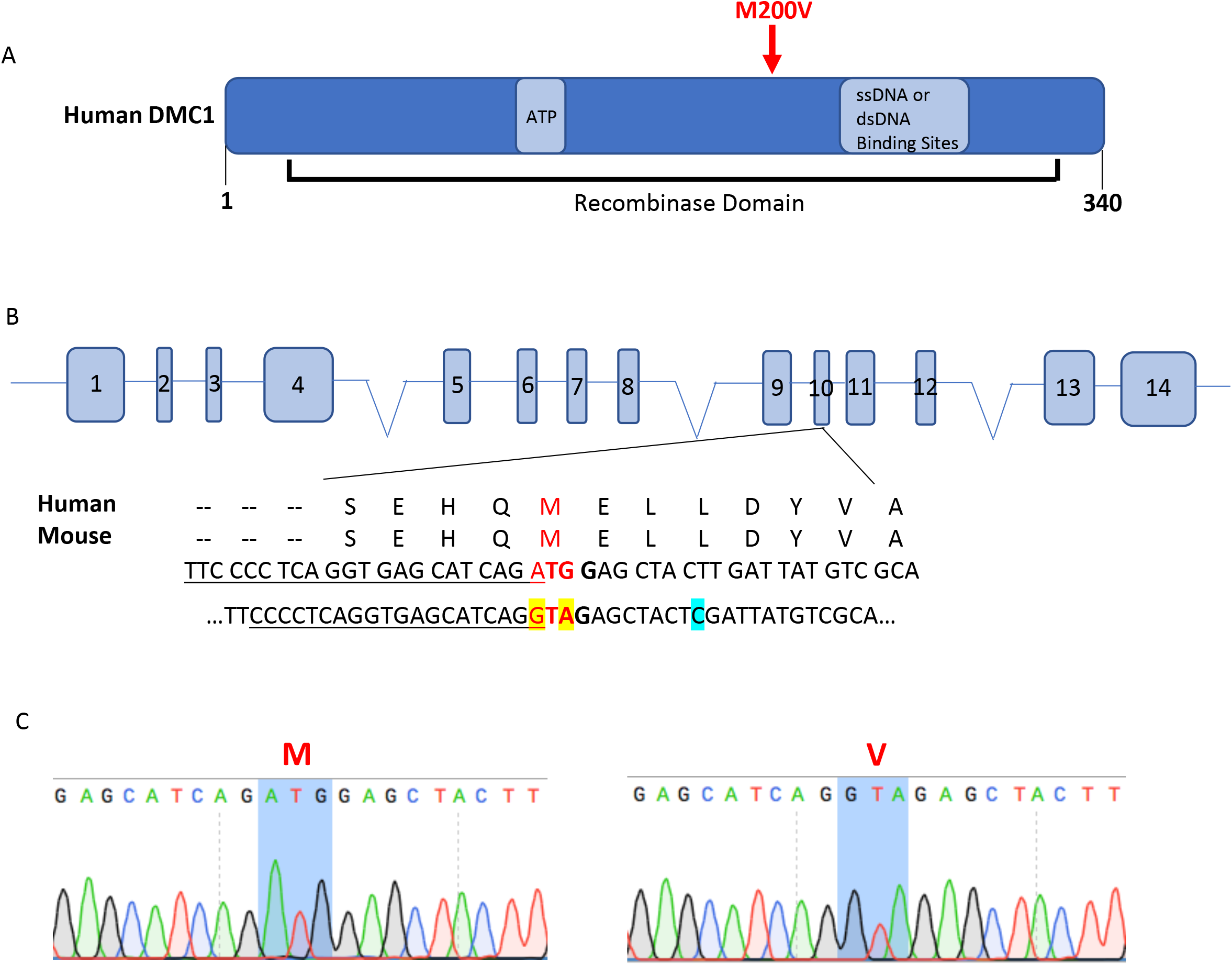
CRISPR/Cas9-mediated generation of *Dmc1^M200V^* mice. A) Diagram of human DMC1 protein labeled with known functional domains and binding sites. Approximate location of amino acid change, M200V (rs2227914), is indicated in red. B) CRISPR-Cas9 genome editing strategy to introduce the M200V amino acid change. The Met200 site is conserved between humans and mice. Underlined is the sgRNA sequence, in bold is the PAM site, highlighted in yellow are nucleotide changes for Val, and highlighted in blue is a silent mutation to introduce a Taq*α*I restriction enzyme site. C) Sanger sequencing chromatograms from wild type (left) and *Dmc1*^M200V/M200V^ mouse (right).

Male and female *Dmc1^M200V/M200V^* mice did not show gross phenotypic abnormalities. To assess the fertility status of these mice, eight-week old *Dmc1^M200V/M200V^* male and female mice were housed with wild type mates over the course of four to eight months. Litter sizes of these mutants vs controls were not significantly different (Figure 2A). Thus, the *Dmc1^M200V^* allele does not impair fertility or fecundity in mice.

**Figure 2.**
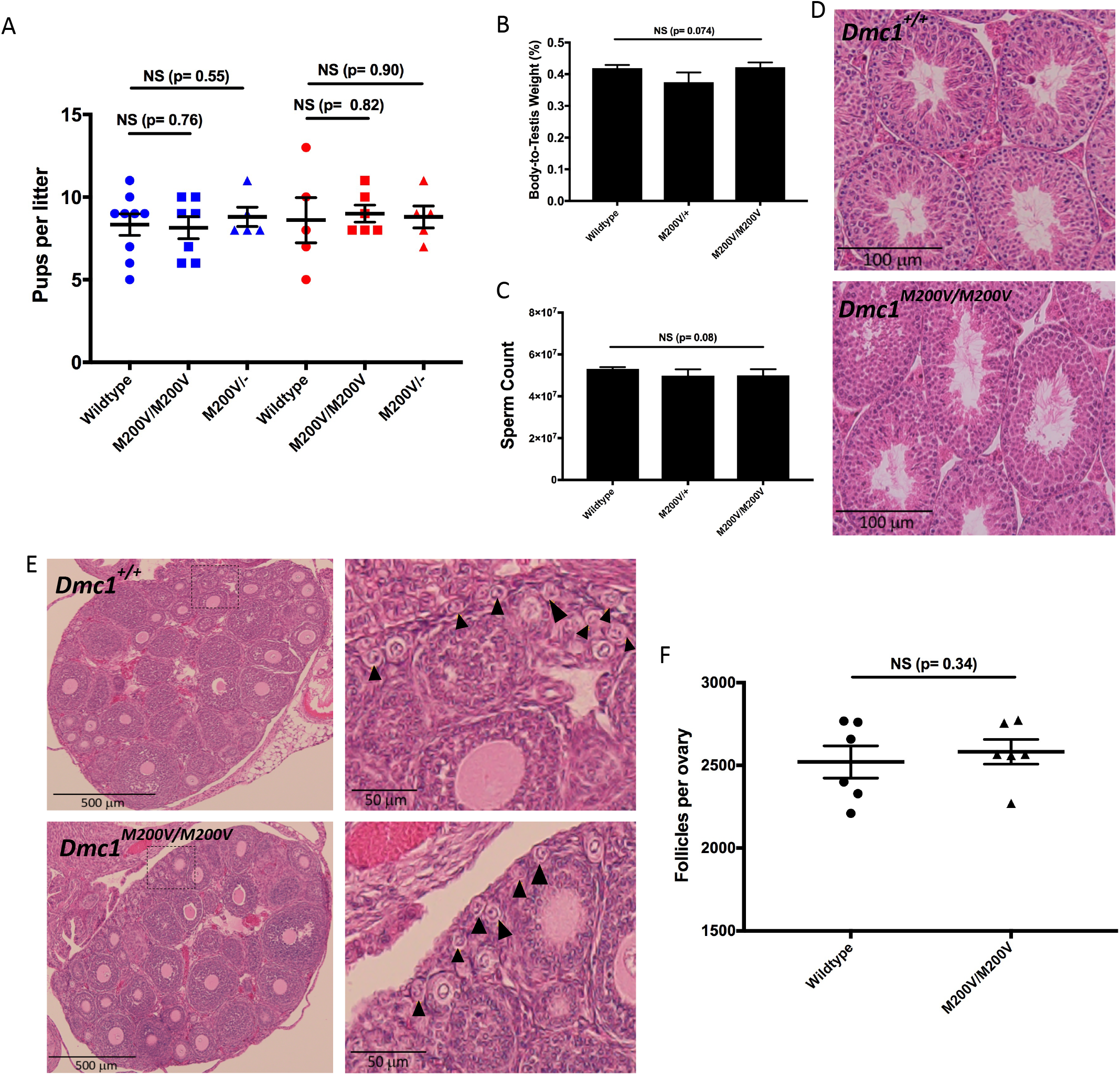
*Dmc1^M200V/+^* and *Dmc1^M200V/M200V^* mice are fertile and show no gross abnormalities. A) Results from fertility test breeding, where each genotype listed was bred to a wild type mate starting at eight-weeks of age. Litter sizes are shown. The first three genotypes on the left (blue) refers to the father’s genotype and the three genotypes on the right (red) refers to the mother’s genotype. We found male:female pups born fit expected Mendelian ratios. Each genotype is n=3 except *Dmc1^M200V/-^* is n=2. Student T-test comparing wild type and *Dmc1^M200V/M200V^* male and female birth rate are p= 0.76 and p= 0.82, respectively. Student T-test comparing wild type and *Dmc1^M200V/-^* male and female birth rate are p= 0.55 and p= 0.90, respectively. B) The mass of eight-week old testes were measured and averaged, then the ratio was calculated relative to the respective animal’s body mass (NS, p=0.074). Wild type n=4, *Dmc1^M200V/+^* n=3, and *Dmc1^M200V/M200V^* n=3. C) Sperm count from eight-week old males (NS, p=0.08). Wild type n=4, *Dmc1^M200V/+^* n=3, and *Dmc1^M200V/M200V^* n=3. D) Representative image of paraffin embedded and H&E stained testis cross-section from eight-week old adult *Dmc1^+/+^* and *Dmc1^M200V/M200V^* at 20x magnification. E) Paraffin-embedded and H&E stained cross-section from three-week old wild type and *Dmc1^M200V/M200V^* ovaries. Magnification starts at 10x. Black arrows mark primordial follicles. F) Quantification of primordial follicles in three-week old ovaries. Averages for each ovary are 2516 follicles in wild type and 2581 follicles in *Dmc1^M200V/M200V^* (NS, p=0.34). Wild type n=3 and *Dmc1^M200V/M200V^* n=3.

To determine if the mutation caused subclinical gonadal defects, gross and histological analyses of testes and ovaries were performed. Eight-week old *Dmc1^M200V/+^* and *Dmc1^M200V/M200V^* testis weights and sperm counts were indistinguishable from wild type (Figure 2B, C). Similarly, mutant testis histology revealed no abnormalities in spermatogenesis (Figure 2D).

Although females had normal fecundity, it is possible that the mutation caused reduction in the ovarian reserve. Wild type and *Dmc1^M200V/M200V^* ovaries were histologically indistinguishable (Figure 2E). We serially sectioned mutant and wild type ovaries and quantified the total number of primordial follicles in three-week old ovaries, revealing no significant difference (Figure 2F). These results are consistent with the breeding studies and provide no evidence of any compromise to gametogenesis in either sex.

### DMC1^M200V^ does not disrupt meiotic chromosome behavior in mice

Despite having normal fertility, fecundity, and histology, it is possible that the M200V amino acid change has a mild effect only detectable at the cellular level. *Dmc1* is a meiosis-specific recombinase that acts at sites of meiotically-programmed double-strand breaks (DSBs) to catalyze D-loop formation and strand exchange between chromosomes, ultimately promoting chromosome pairing, synapsis, and recombination (Yoshida et al. 1998; Bishop et al. 1992; Schwacha and Kleckner 1997; D. L. Pittman et al. 1998). As mentioned in the Introduction, in vitro assays showed that DMC1^M200V^ had altered biochemical activities under certain conditions, namely reduced ATPase activity at higher temperatures and reduced D-loop formation at low magnesium concentrations. Furthermore, the orthologous amino acid change in the fission yeast *S. pombe* caused a 50% reduction in recombination at two loci (Hikiba et al. 2008). To detect any anomalies that may reflect defective function of DMC1^M200V^ during mouse meiosis, we conducted immunocytological analyses of surface-spread chromosomes from wild type and *Dmc1^M200V/M200V^* spermatocytes, using markers diagnostic for certain key events of meiosis. First, we tested if DMC1^M200V^ localizes normally to meiotically-programmed DSBs. There was no significant difference in DMC1 focus numbers between wild type and *Dmc1^M200V/M200V^* during the peak stages of DSB formation: leptonema and zygonema (Figure 3A, B). Normally, as meiotic prophase I progresses and DSBs are repaired by recombination, DSBs as marked by DMC1 foci along the lateral/axial elements of synaptonemal complexes (marked by SYCP3) decrease in zygonema and pachynema, ultimately disappearing in late pachynema. This was the case with *Dmc1^M200V/M200V^* spermatocytes. These results indicate that DMC1^M200V^ is properly recruited to sites of meiotic DSBs, catalyzes homologous recombination repair of these DSBs which is essential for pairing of homologous chromosomes, and is properly released from processed DSBs following repair.

**Figure 3.**
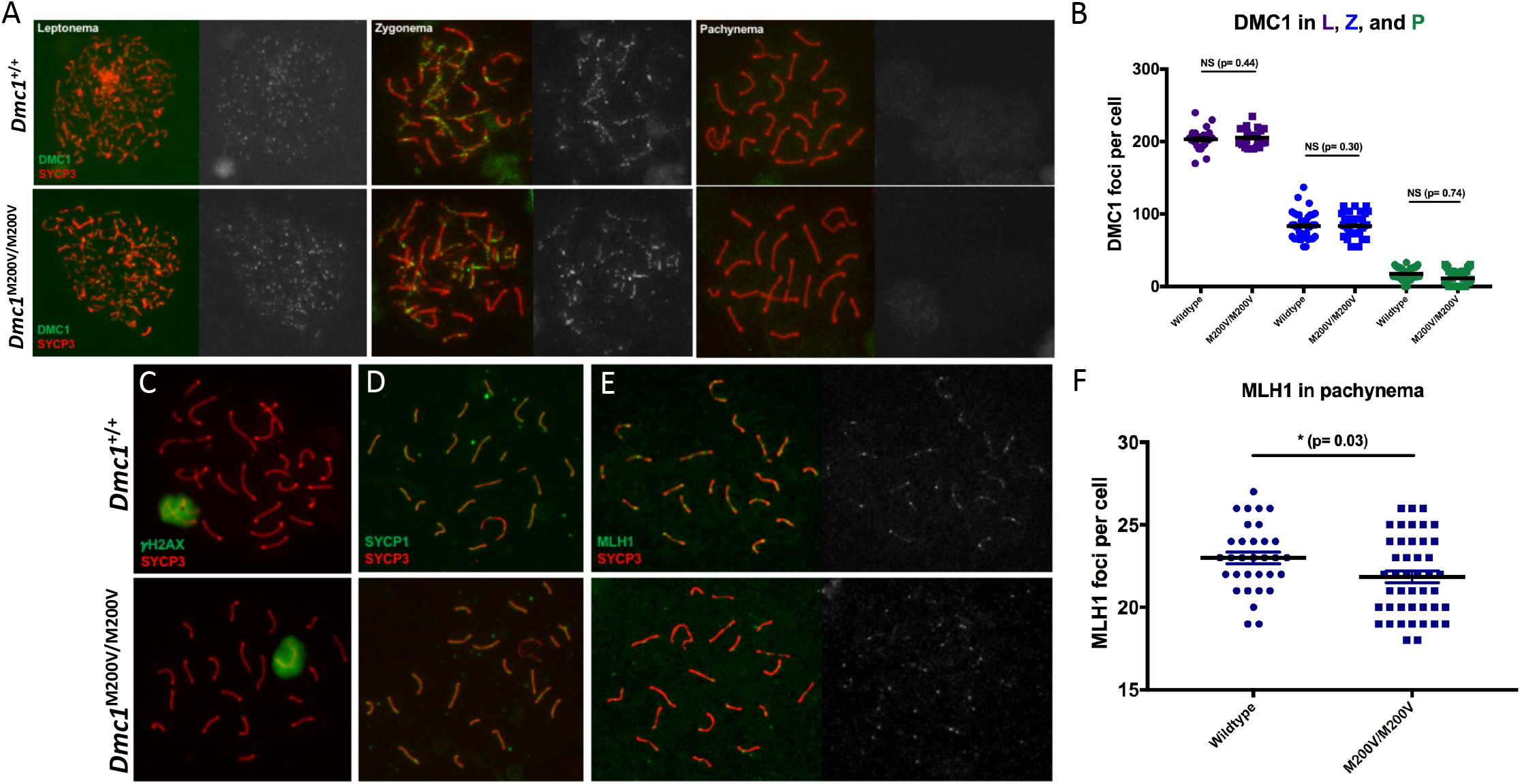
Chromosome surface-spreads show normal recruitment and initiation of recombination in *Dmc1^M200V/M200V^* spermatocytes. A) From left to right are spreads of leptotene, zygotene, and pachytene spermatocyte nuclei stained for SYCP3 (red; which marks axial/lateral elements of forming or formed synapstonemal complexes) and DMC1 (green). B) Quantification of DMC1 foci at leptonema, zygonema, and pachynema. Average DMC1 foci in wild type is 206 foci/cell at leptonema (25 cells), 79 foci/cell at zygonema (30 cells), and 16 foci/cell in pachynema (30 cells). The average DMC1 foci in *Dmc1^M200V/M200V^* is 208 foci/cell in leptonema (45 cells, NS, p=0.44), 83 foci/cell in zygonema (40 cells, NS, p=0.30), and 11 foci/cell in pachynema (38 cells, NS, p=0.74). Wild type n=3 and *Dmc1^M200V/M200V^* n=3. C) Representative spreads of pachytene cells stained for *γ*H2AX (green) and SYCP3 (red). Note normal XY body appearance in *Dmc1^M200V/M200V^* nuclei. Wild type n=3 and *Dmc1^M200V/M200V^* n=3. D) Representative spreads of pachytene cells stained for synaptonemal complex proteins, SYCP1 (green) and SYCP3 (red). *Dmc1^M200V/M200V^* achieves full synapsis. Wild type n=3 and *Dmc1^M200V/M200V^* n=3. E) Representative spreads of pachytene cells stained for MLH1 (green) and SYCP3 (red). F) Quantification of MLH1 foci in pachynema. Average MLH1 foci in wild type is 23±2.0 foci/cell (n=3, 96 cells) and *Dmc1^M200V/M200V^* is 22±2.4 foci/cell (n=4, 120 cells). There is statistical significance using two-tailed student T-test, p=0.03, between wild type and *Dmc1^M200V/M200V^*.

Next, we immunolabeled spreads with *γ*H2AX, a histone variant phosphorylated at sites of DNA damage, which also serves as a marker of the transcriptionally silenced and heterochromatinized XY body that forms in pachynema (Burma et al. 2001). Similar to wild type, leptotene and zygotene *Dmc1^M200V/M200V^* spermatocytes had high levels of *γ*H2AX along chromosomes (data not shown) and by pachynema, *γ*H2AX was exclusive to the X and Y chromosomes, indicative of complete DSB repair on autosomes and XY body formation (Figure 3C). We then immunolabeled chromosome spreads for synaptonemal complex (SC) transverse element protein SYCP1, which marks regions of complete synapsis between homologs. The former decorates the axial/lateral elements of the synaptonemal complex, while the latter is a component of the transverse element, and thus only stains synapsed regions of chromosomes. In the absence of DMC1, repair of meiotic DSBs does not occur, and homologous chromosomes do not synapse (D. Pittman et al. 1998; Yoshida et al. 1998). SC formation occurred normally in *Dmc1*^M200V/M200V^ spermatocytes as indicated by SYCP3/SYCP1 staining of pachytene chromosomes (Figure 3D). This suggests that DMC1^M200V^ does not affect homologous chromosome synapsis.

Meiotic DSBs are repaired by one of two mechanisms: the majority (~90%) as noncrossovers (NCOs) and the rest as crossovers (COs). While both types of events are important for driving homolog pairing and synapsis, chiasmata formed by CO events are essential for proper chromosome segregation at the first meiotic division. They ensure that homologs correctly align at the metaphase plate and segregate to opposite poles when the COs are resolved. To test whether crossover formation is affected in *Dmc1^M200V/M200V^* mutants, we quantified MLH1 foci - a proxy for the predominant class of crossovers in mice - in pachytene spermatocyte nuclei (Figure 3E). Interestingly, whereas wild type spermatocytes had 23 ± 2.0 MLH1 foci/cell (mean±s.d., n=3, 96 cells), *Dmc1^M200V/M200V^* had 22 ± 2.4 MLH1 foci/cell (mean±s.d., n=4, 120 cells), which is a small but statistically significant difference (p= 0.03; Figure 3F).

### One copy of *Dmc1^M200V^* is sufficient for normal meiosis

Although fertility, fecundity, and meiotic prophase I chromosome behavior was normal in *Dmc1^M200V/M200V^* mice, we considered the possibility that the mutant protein might be compromised in a subtle way that would only manifest under more challenging conditions. Accordingly, we bred mice hemizygous for *Dmc1^M200V^* (*Dmc1^M200V/-^*). The testis weights and sperm counts of eight-week old *Dmc1^M200V/-^* males were no different than wild type (Figure 4A,B), and testis histology was unremarkable (Figure 4C). Likewise, females at three weeks of age had primordial follicle counts similar to wildtype (Figure 4D). Lastly, breeding *Dmc1^M200V/-^* animals to wildtype mates yielded normal sized litters; males sired an average of 8.8 pups/litter and females birthed 8.8 pups/litter (p= 0.55 and p= 0.90, respectively; Figure 2A). These results provide further evidence that under *in vivo* conditions, DMC1^M200V^ has no substantial impact on fertility, fecundity, or gametogenesis.

**Figure 4.**
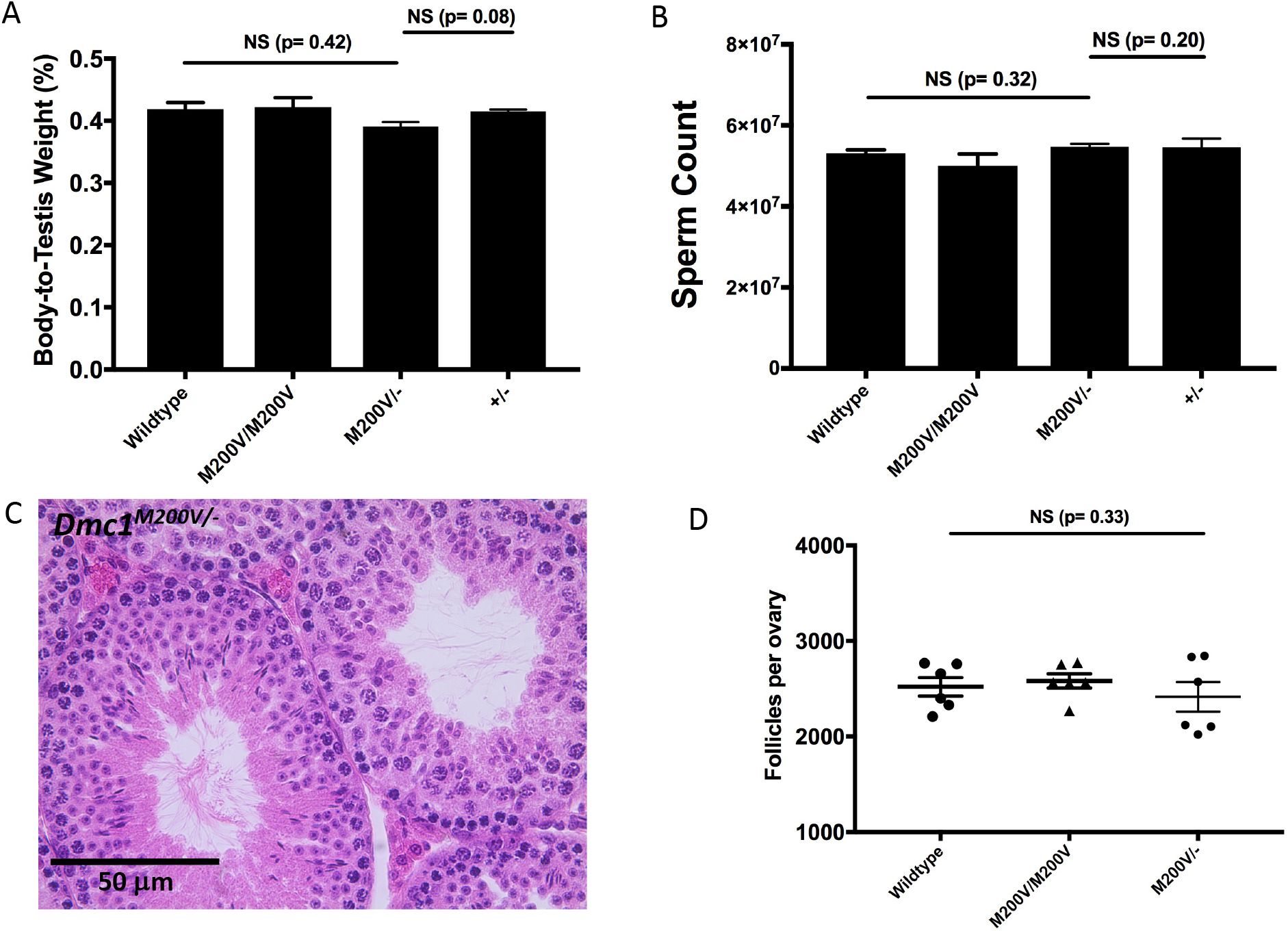
One copy of *Dmc1^M200V^* allele is sufficient for normal meiosis and fertility in mice. A) Data from Figure 2B plus testis sizes from eight-week old *Dmc1^M200V/-^* (n=4) and *Dmc1*^+/-^ (n=2) males. Student T-test between wild type and *Dmc1^M200V/-^* is p=0.42. B) Data from Figure 2C plus epididymis sperm counts from eight-week old *Dmc1^M200V/-^* (n=4) and *Dmc1^+/-^* (n=2) males. Student T-test between wild type and *Dmc1^M200V/-^* is p=0.32. C) Representative image of paraffin-embedded and H&E stained testis cross-section from eight-week old *Dmc1^M200V/-^* male. D) Data from Figure 2F plus primordial follicle counts from three-week old *Dmc1^M200V/-^* ovaries (average=2416 follicles/ovary, p=0.33, n=3).

## DISCUSSION

Remarkable advances in genomics capabilities in the past few years are revolutionizing medicine. As DNA sequencing costs have dropped precipitously, it is becoming more routine to use whole-exome or -genome sequencing as clinical tests for certain common (namely, cancer) or rare diseases (such as undiagnosed developmental disorders). This will certainly become more commonplace for identifying potential genetic causes of idiopathic infertility in individual patients. Coupled with the advent of genome editing technology, it is increasingly plausible (scientifically) to envision genetic correction of infertility-causing alleles, particularly in males where spermatogonial stem cells are present and can be cultured, manipulated, and returned into the patient. Of course such interventions, should they indeed be deemed as safe and acceptable, are absolutely dependent upon unequivocal identification of the infertility-causing mutation or variant.

The work presented here is part of an NIH-funded project (see acknowledgements) we have undertaken to specifically address this issue (Singh and Schimenti 2015), and our results serve to highlight the serious nature of the problem. The *Dmc1^M200V^* allele was identified as a potentially causative infertility variant 10 years ago (Mandon-Pépin et al. 2008; Hikiba et al. 2008) and associated work provided compelling evidence that the allele encoded a protein with altered biochemical activities that might be consistent with disrupted meiosis (Hikiba et al. 2008). However, our in vivo modeling of this allele showed that this allele, in mice, did not impact fertility or fecundity at all.

While we are of the opinion that *in vivo* modeling in mice is a more physiologically-relevant biological test than *in vitro* biochemical assays, there are still potential caveats. One caveat is that although the mouse and human DMC1 proteins are 97% identical, it is possible that the M200V alteration is tolerated in mice but not humans. We consider this unlikely because there are no *Dmc1* paralogs unique to mice, and the mouse gene is absolutely required for meiosis and fertility in mice. It is also conceivable that the gonadal environment in humans is such that the M200V change is catalytically more unstable in human meiocytes than in mouse meiocytes. For example, we found that in *Dmc1^M200V/M200V^* spermatocytes, there was a slight decrease (<5%) in the number of MLH1 foci. However, the number of MLH1 foci varies amongst inbred strains (Anderson et al. 1999), and since the founder mice were of mixed genetic background, it is possible that the variation is due to these differences, possibly linked to the *Dmc1* locus. Nevertheless, the impact, if any, was negligible in terms of all other reproductive parameters (e.g. germ cell numbers, litter sizes, etc.).

The case of *Dmc1^M200V^* is emblematic of the challenges facing genetic diagnosis of infertility in a clinical situation. Computational algorithms are useful for estimating the potential damage of a variant on protein function, but should not serve as an endpoint during the validation process due to their insufficient reliability (Wang et al. 2018; Miosge et al. 2015), particularly if the information was to be used for treatment actions. The DMC1^M200V^ allele was originally cited as being “probably damaging” by the widely-used Polyphen algorithm (Mandon-Pépin et al. 2008). However, the updated Polyphen-2 now predicts the variant as benign, as does FATHMM, SNPs & GO, and SIFT. In contrast, PANTHER and Mutation Assessor score the allele as being “possibly damaging” and “medium functional impact”, respectively. Current algorithms measure different criteria with different emphases, which can result in contradictory conclusions. However, technology is dynamic, and with increasing numbers of *in vivo* studies and training data sets, these algorithms may become more accurate with time. Another indicator of whether an allele might cause infertility is allele frequency. As noted by He *et al.* (He et al. 2018), the *Dmc1^M200V^* has a high allelic frequency in African populations (12% according to the gnomAD database), which seems incompatible with it being an infertility allele.

In conclusion, this study highlights the importance of *in vivo* modeling to make accurate conclusions about putative infertility-associated polymorphisms or *de novo* mutations before taking clinical actions. While mice are currently the most accurate models for this (Singh and Schimenti 2015; Fujihara et al. 2017; Kherraf et al. 2018), we anticipate that improvements to *in vitro* gametogenesis, especially with human cells, will be important for development of higher-throughput and lower-cost analysis for diagnostics of genetic causes of human infertility.

## Acknowledgements

We would like to thank Robert Munroe and Christian Abratte at Cornell University’s Stem Cell and Transgenic Core Facility (supported by contract C029155 from the New York State Stem Cell program) for performing the CRISPR-Cas9 microinjections. This work was also supported by a grant from the National Institute of Child Health and Human Development (R01HD082568).

## Materials and Methods

### Generation of *Dmc1^M200V^* Mice by CRISPR-Cas9 Genome Editing

An optimal guide sequence was selected using online software at mit.crispr.edu. The crRNA and CRISPR-Cas9 tracrRNA was synthesized by IDT (ALT-R service) and the ssODN was also synthesized by IDT’s (Ultramer service). Prior to pronuclear injection, the crRNA (25 ng/μL) and tracrRNA (25 ng/μL) were co-incubated to form a ribonucleoprotein complex according to manufacturer’s instructions. The ssODN (50 ng/μL) and additional CAS9 protein (1000 ng/μL; PNA Bio) were added, and all materials were co-injected into zygotes (F1 hybrids between strains FVB/NJ and B6(Cg)-Tyr^c-2J^/J), then transferred into the oviduct of pseudopregnant females. Founders bearing at least one copy of the desired alteration were identified by PCR with primers flanking the SNP (Supp. Table I), then backcrossed into FVB/NJ. Initial phenotyping was performed after one backcross generation followed by intercrossing, then additional phenotyping was done with animals backcrossed an additional two or more generations. No phenotypic differences were found among different generations.

All animal use was conducted under a protocol (2004-0038) to J.C.S. and approved by Cornell University’s Institutional Animal Use and Care Committee.

The *Dmc1^tm1Jcs^* allele (referred to as *Dmc1^+/-^* in this manuscript) was previously described (Pittman et al. 1998).

All sequences used for CRISPR-Cas9 editing and mouse genotyping are in Supplementary Table I.

### Genotyping of *Dmc1^M200V^* and *Dmc1^+/-^* Mice

Toe tissue was collected from pups at 8-14 days of age. A crude DNA lysate was made by disrupting tissue in Lysis Buffer (2.5 mM NaOH/0.2 mM EDTA, pH 12.2) at 95°C for 60 minutes and neutralized by adding equal parts Neutralization Buffer (40 mM Tris-HCl, pH 4.6). PCR was done using Econotaq and associated reagents (Lucigen), following manufacturer’s protocol with 3 μL of crude DNA lysate per reaction. Primers were ordered from IDT and sequences are listed in Supplementary Table I. The PCR cycle used for both *Dmc1^M200V^* and *Dmc1^+/-^* mice PCR was: initial denature at 95°C for 5 minutes, 30 cycles of 95°C for 30 seconds - 58°C for 30 seconds - 72°C for 30 seconds, and final elongation at 72°C for 5 minutes. For identification of *Dmc1^+/-^* mice, we used gel electrophoresis to analyze the presence of wildtype (167bp) and/or mutant PCR (250bp) products amplified by respective primers. For identification of *Dmc1^M200V^* mice, PCR samples were digested with restriction enzyme Taq*α*I (NEB) at 65°C for two hours then analyzed using gel electrophoresis. The wildtype allele is detected as two bands (444bp and 31bp) while the *Dmc1^M200V^* allele is detected as three bands (244bp, 200bp, and 31bp).

### Histology and Primordial Follicle Quantification

Testes were harvested from eight-week old males and fixed in Bouin’s for 24 hours, washed in 70% ethanol for 24 hours, and embedded in paraffin. Sections of 6 μm were made and stained with hematoxylin and eosin. Ovaries were harvested from three week old females and prepared in the same way, then serial sectioned at 6 μm. Primordial follicles were counted in every fifth section and final follicle counts were calculated as previously described (Myers et al. 2004). Statistical analysis was done with a two-tailed student T-test on Prism 7 (GraphPad).

### Sperm Counts

Epidydymides were isolated from eight-week old males. The tissue was minced in 5mL of MEM media and incubated at 37°C for 15 minutes, allowing spermatozoa to swim out. The spermatozoa were diluted 1:2 in MEM and counted using a hemacytometer.

### Immunocytochemistry of Meiotic Chromosomes

We used a protocol that has been previously published (McNairn et al. 2017). In brief, testes were isolated from eight-to twelve-week old males, detunicated, and minced in MEM media. Spermatocytes were hypotonically swollen in 4.5% sucrose solution, lysed in 0.1% Triton X-100, 0.02% SDS, and 2% formalin. Slides were washed and stained immediately or stored at -80°C. Blocking buffer used was 5% goat serum in PBS, 0.1% Tween20 and slides were blocked for 1 hour at room temperature. Primary antibodies and dilutions used were: anti-SYCP3 (1:600, Abcam, #ab15093), anti-SYCP3 (1:600, Abcam, #ab97672), anti-SYCP1 (1:400, Abcam, #ab15090), anti-DMC1 (1:100, Abcam, #ab11054), anti-MLH1 (1:100, BD Pharmingen, #554073), and anti-phospho-H2A.X (1:1000, Millipore, #16-193). Primary staining of chromosome surface spreads was done at 37°C and incubated overnight. Secondary antibodies used were: goat anti-mouse IgG AlexaFluor 488 (1:1000, ThermoFisher Scientific, #R37120) and goat anti-rabbit IgG AlexaFluor 594 (1:800, ThermoFisher Scientific, #R37117). Secondary antibodies were incubated for 1 hour at room temperature. Images were acquired with an Olympus microscope with 40x lens using cellSens software (Olympus). Foci were quantified using ImageJ with plugins Cell Counter (Kurt De Vos) and Nucleus Counter. All data was analyzed in Prism 7 (GraphPad).

---

**Supplementary Table I.**
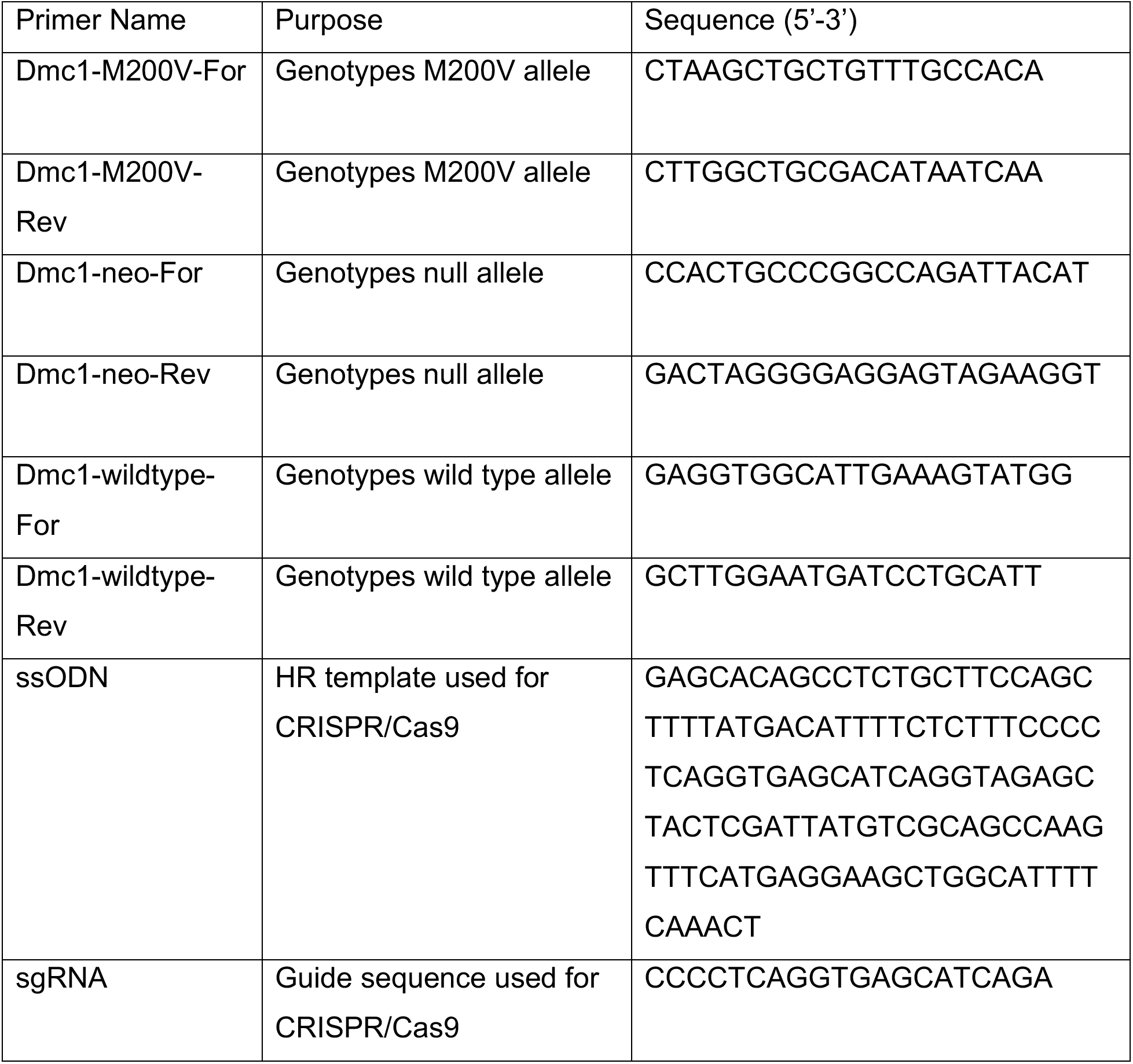

## BIBLIOGRAPHY

Adzhubei, I.A., Schmidt, S., Peshkin, L., Ramensky, V.E., Gerasimova, A., Bork, P., Kondrashov, A.S. and Sunyaev, S.R. 2010. A method and server for predicting damaging missense mutations. Nature Methods 7(4), pp. 248–249.

Anderson, L.K., Reeves, A., Webb, L.M. and Ashley, T. 1999. Distribution of crossing over on mouse synaptonemal complexes using immunofluorescent localization of MLH1 protein. Genetics 151(4), pp. 1569–1579.

Bishop, D.K., Park, D., Xu, L. and Kleckner, N. 1992. DMC1: a meiosis-specific yeast homolog of E. coli recA required for recombination, synaptonemal complex formation, and cell cycle progression. Cell 69(3), pp. 439–456.

Bolcun-Filas, E., Rinaldi, V.D., White, M.E. and Schimenti, J.C. 2014. Reversal of female infertility by Chk2 ablation reveals the oocyte DNA damage checkpoint pathway. Science 343(6170), pp. 533–536.

Burma, S., Chen, B.P., Murphy, M., Kurimasa, A. and Chen, D.J. 2001. ATM phosphorylates histone H2AX in response to DNA double-strand breaks. The Journal of Biological Chemistry 276(45), pp. 42462–42467.

Dam, A.H.D.M., Koscinski, I., Kremer, J.A.M., Moutou, C., Jaeger, A.-S., Oudakker, A.R., Tournaye, H., Charlet, N., Lagier-Tourenne, C., van Bokhoven, H. and Viville, S. 2007. Homozygous mutation in SPATA16 is associated with male infertility in human globozoospermia. American Journal of Human Genetics 81(4), pp. 813–820.

França, M.M., Lerario, A.M., Funari, M.F.A., Nishi, M.Y., Narcizo, A.M., de Mello, M.P., Guerra-Junior, G., Maciel-Guerra, A.T. and Mendonça, B.B. 2017. A Novel Homozygous Missense FSHR Variant Associated with Hypergonadotropic Hypogonadism in Two Siblings from a Brazilian Family. Sexual Development: Genetics, Molecular Biology, Evolution, Endocrinology, Embryology, and Pathology of Sex Determination and Differentiation 11(3), pp. 137–142.

Fujihara, Y., Oji, A., Larasati, T., Kojima-Kita, K. and Ikawa, M. 2017. Human Globozoospermia-Related Gene Spata16 Is Required for Sperm Formation Revealed by CRISPR/Cas9-Mediated Mouse Models. International Journal of Molecular Sciences 18(10).

He, W.-B., Tu, C.-F., Liu, Q., Meng, L.-L., Yuan, S.-M., Luo, A.-X., He, F.-S., Shen, J., Li, W., Du, J., Zhong, C.-G., Lu, G.-X., Lin, G., Fan, L.-Q. and Tan, Y.-Q. 2018. DMC1 mutation that causes human non-obstructive azoospermia and premature ovarian insufficiency identified by whole-exome sequencing. Journal of Medical Genetics 55(3), pp. 198–204.

Hikiba, J., Hirota, K., Kagawa, W., Ikawa, S., Kinebuchi, T., Sakane, I., Takizawa, Y., Yokoyama, S., Mandon-Pépin, B., Nicolas, A., Shibata, T., Ohta, K. and Kurumizaka, H. 2008. Structural and functional analyses of the DMC1-M200V polymorphism found in the human population. Nucleic Acids Research 36(12), pp. 4181–4190.

Keeney, S., Giroux, C.N. and Kleckner, N. 1997. Meiosis-specific DNA double-strand breaks are catalyzed by Spo11, a member of a widely conserved protein family. Cell 88(3), pp. 375–384.

Kherraf, Z.-E., Conne, B., Amiri-Yekta, A., Kent, M.C., Coutton, C., Escoffier, J., Nef, S., Arnoult, C. and Ray, P.F. 2018. Creation of knock out and knock in mice by CRISPR/Cas9 to validate candidate genes for human male infertility, interest, difficulties and feasibility. Molecular and Cellular Endocrinology.

Kumar, P., Henikoff, S. and Ng, P.C. 2009. Predicting the effects of coding non-synonymous variants on protein function using the SIFT algorithm. Nature Protocols 4(7), pp. 1073–1081.

Mandon-Pépin, B., Touraine, P., Kuttenn, F., Derbois, C., Rouxel, A., Matsuda, F., Nicolas, A., Cotinot, C. and Fellous, M. 2008. Genetic investigation of four meiotic genes in women with premature ovarian failure. European Journal of Endocrinology 158(1), pp. 107–115.

McNairn, A.J., Rinaldi, V.D. and Schimenti, J.C. 2017. Repair of meiotic DNA breaks and homolog pairing in mouse meiosis requires a minichromosome maintenance (MCM) paralog. Genetics 205(2), pp. 529–537.

Miosge, L.A., Field, M.A., Sontani, Y., Cho, V., Johnson, S., Palkova, A., Balakishnan, B., Liang, R., Zhang, Y., Lyon, S., Beutler, B., Whittle, B., Bertram, E.M., Enders, A., Goodnow, C.C. and Andrews, T.D. 2015. Comparison of predicted and actual consequences of missense mutations. Proceedings of the National Academy of Sciences of the United States of America 112(37), pp. E5189–98.

Miyamoto, T., Koh, E., Sakugawa, N., Sato, H., Hayashi, H., Namiki, M. and Sengoku, K. 2008. Two single nucleotide polymorphisms in PRDM9 (MEISETZ) gene may be a genetic risk factor for Japanese patients with azoospermia by meiotic arrest. Journal of Assisted Reproduction and Genetics 25(11–12), pp. 553–557.

Miyamoto, T., Tsujimura, A., Miyagawa, Y., Koh, E., Sakugawa, N., Miyakawa, H., Sato, H., Namiki, M., Okuyama, A. and Sengoku, K. 2009. A single nucleotide polymorphism in SPATA17 may be a genetic risk factor for Japanese patients with meiotic arrest. Asian Journal of Andrology 11(5), pp. 623–628.

Myers, M., Britt, K.L., Wreford, N.G.M., Ebling, F.J.P. and Kerr, J.B. 2004. Methods for quantifying follicular numbers within the mouse ovary. Reproduction 127(5), pp. 569–580.

Pittman, D., Cobb, J., Schimenti, K., Wilson, L., Cooper, D., Brignull, E., Handel, M.A. and Schimenti, J. 1998. Meiotic prophase arrest with failure of chromosome pairing and synapsis in mice deficient for. Mol. Cell 1, pp. 697–705.

Pittman, D.L., Cobb, J., Schimenti, K.J., Wilson, L.A., Cooper, D.M., Brignull, E., Handel, M.A. and Schimenti, J.C. 1998. Meiotic prophase arrest with failure of chromosome synapsis in mice deficient for Dmc1, a germline-specific RecA homolog. Molecular Cell 1(5), pp. 697–705.

Robert, T., Nore, A., Brun, C., Maffre, C., Crimi, B., Bourbon, H.M. and de Massy, B. 2016. The TopoVIB-Like protein family is required for meiotic DNA double-strand break formation. Science 351(6276), pp. 943–949.

Sato, H., Miyamoto, T., Yogev, L., Namiki, M., Koh, E., Hayashi, H., Sasaki, Y., Ishikawa, M., Lamb, D.J., Matsumoto, N., Birk, O.S., Niikawa, N. and Sengoku, K. 2006. Polymorphic alleles of the human MEI1 gene are associated with human azoospermia by meiotic arrest. Journal of Human Genetics 51(6), pp. 533–540.

Schwacha, A. and Kleckner, N. 1997. Interhomolog bias during meiotic recombination: meiotic functions promote a highly differentiated interhomolog-only pathway. Cell 90(6), pp. 1123–1135.

Shihab, H.A., Gough, J., Cooper, D.N., Stenson, P.D., Barker, G.L.A., Edwards, K.J., Day, I.N.M. and Gaunt, T.R. 2013. Predicting the functional, molecular, and phenotypic consequences of amino acid substitutions using hidden Markov models. Human Mutation 34(1), pp. 57–65.

Singh, P. and Schimenti, J.C. 2015. The genetics of human infertility by functional interrogation of SNPs in mice. Proceedings of the National Academy of Sciences of the United States of America 112(33), pp. 10431–10436.

Wang, T., Bu, C.H., Hildebrand, S., Jia, G., Siggs, O.M., Lyon, S., Pratt, D., Scott, L., Russell, J., Ludwig, S., Murray, A.R., Moresco, E.M.Y. and Beutler, B. 2018. Probability of phenotypically detectable protein damage by ENU-induced mutations in the Mutagenetix database. Nature Communications 9(1), p. 441.

Yoshida, K., Kondoh, G., Matsuda, Y., Habu, T., Nishimune, Y. and Morita, T. 1998. The mouse RecA-like gene Dmc1 is required for homologous chromosome synapsis during meiosis. Molecular Cell 1(5), pp. 707–718.

